# Estimating haplotype values and mutation effects in the context of a local DNA tree

**DOI:** 10.1101/2025.11.08.687335

**Authors:** Gabriela Mafra Fortuna, Jana Obšteter, Ajda Moškrič, Gregor Gorjanc

## Abstract

**Background:** Single nucleotide polymorphism (SNP) arrays provide genome-wide coverage of polymorphic sites across populations, but most quantitative genetics models assume that the effects of these mutations remain constant across generations and populations. This assumption overlooks population dynamics and genetic architecture that are central to trait expression, particularly when inferences are made across populations. Whole-genome sequence (WGS) data captures all variants and should, in principle, overcome these limitations, but its use has delivered only modest gains in prediction accuracy at considerable computational cost. Ancestral recombination graphs (ARGs) offer a representation of genome variations that describes how genetic variation is shaped by haplotype inheritance between generations (represented as local DNA trees) and associated mutations. This study investigates how a generative model on a local DNA tree can improve the estimation of mutation and haplotype effects, especially for rare or population-specific variants.

**Methods:** We developed a TBLUP approach that uses local DNA tree information to estimate haplotype and mutation effects. In each local DNA tree, branches connecting ancestral and descendant haplotypes represent DNA inheritance, and the trait associated with a branch corresponds to the mutation(s) it carries. Summing these branch-specific mutation effects from the tree root to each haplotype defines the haplotype values. This recursive structure yields a sparse and computationally efficient approach for estimating haplotype and mutation effects from the local DNA tree and phenotypes. We show how the TBLUP approach is similar and different to the SNP-BLUP and GBLUP approaches and demonstrate it with cattle mitochondrial DNA, a non-recombining genomic region, using both simulated and real data.

**Results and conclusions:** The TBLUP approach was computationally more efficient than SNP-BLUP/GBLUP approaches and produced more accurate estimates of haplotype values and mutation effects, which can vary between haplotypes. The accuracy increased when phenotypes were available for haplotypes across the local DNA tree rather than only for the recent haplotypes. Incorporating local DNA tree information enhances the use of genomic data in quantitative genetics. Extending the TBLUP approach to full ARGs will enable analysis across multiple local DNA trees (accounting and leveraging recombination), which will further improve quantitative genetic modelling and practical applications.

## Background

Genomic selection has revolutionised animal breeding. The use of a large number of genome-wide SNP markers combined with powerful statistical models has enabled more informed quantitative genetic analyses and more accurate or earlier breeding decisions compared to traditional pedigree-based approaches [1, 2, 3]. One of the most successful implementations of genomic selection has been in dairy cattle. There, the use of genomic predictions enabled halving the generation interval for the male path of selection and, consequently, doubled the rate of genetic gain per unit of time [4, 5]. Similar success stories have also been reported for the implementation of genomic selection in other species [6, 7].

Despite the undeniable power of genomic data for quantitative genetics and animal breeding applications, challenges and opportunities for further improvement remain. Mutations and their effects at causal loci can differ across genetic distances, such as between generations or populations, yet most quantitative genomic models assume constant effects. Drivers of such differences in mutations and their effects include allelic heterogeneity, dominance and epistatic interaction effects, genotype-by-environment interaction effects, and population genetic processes that generate sufficient allele and genotype frequency variation at sites contributing to these effects [8, 9, 10, 11, 12, 13, 14, 15]. Theoretical modelling and simulations showed how mutation effects change due to interactions and population genetic processes [16, 11, 17, 18, 19], which is corroborated with empirical results from a range of species [20, 21, 22, 23]. The term epistatic drift has been proposed to describe these changes in mutation effects [20]. However, two recent studies of human admixed populations found very high correlation between effect sizes from multiple continental ancestral populations [24, 25]. Perhaps this is due to the fact that the majority of the genetic variance for traits is statistically additive [26] and very large datasets are required to estimate the interaction effects [27, 28].

Another challenge is that most applications use marker genotype data from SNP arrays. Many of these SNP markers were chosen because they are polymorphic across several populations (meaning that they largely predate population splits) and are close to uniformly covering the genome. While this is an effective approach to capture substantial amounts of genetic variation between and within populations, SNP array do not capture many of the rare or recent mutations [29]. Since many causal loci with larger mutation effects are expected to host rare or recent mutations and many causal loci with smaller mutation effects are expected to host older mutations [30], both of these effects are captured through an intricate linkage-disequilibrium between causal and marker loci [1, 31], but also between the loci and population or pedigree structure [32, 33, 34, 35]. Because linkage-disequilibrium is dynamic across generations of a population and across populations, so are estimates of allele substitution effects at the marker loci with respect to the true mutation effects at the causal loci. This dynamics is driven by the genetic distances between genotyped individuals (and the associated time for recombinations between marker and causal loci) [32, 33, 34, 36, 37, 38, 39, 40, 41] and it varies between traits and contexts, also indicating the importance of trait evolution, genetic architecture, and contexts in which data are collected [42, 43].

The challenges with rare mutations and estimating drifting mutation effects via linked SNP markers are particularly relevant for genomic predictions across generations within a population or even across populations, where the genetic distance between training and prediction sets is large. Such genomic predictions are particularly important for numerically small breeding programmes with limited training set sizes [44] and in crossbreeding programs, where it is relevant to predict across populations [45]. The accuracy of these predictions is moderate, although expanding the genomic prediction model into main and population-specific effects has been shown to increase the accuracy somewhat, with mixed results across studies [45, 46, 47, 48, 49, 50, 51, 52]. These observed accuracies mirror the results from human genetics, where transferability of polygenic risk scores is a challenge due to potential differences in genetic architecture across populations, past demography and selection effects, and other factors [41, 53, 54, 55, 56, 57].

Theoretically, some of these challenges could be addressed by replacing SNP array data with whole-genome sequence (WGS). The advantages of WGS would come from capturing more genome variation, avoiding the ascertainment bias of SNP array data, and encompassing both marker and causal loci, which would enable more accurate estimates between generations as well as across larger genetic distances [58, 59, 60, 61]. However, genomic prediction with WGS has shown marginal improvements in accuracy over SNP arrays with inconsistent results between studies (see review from [62]), even with the largest WGS efforts to date [63]. One reason for this is that these studies mostly used biallelic SNPs while ignoring structural variants, and have significantly leveraged imputation to increase the training set size. The benefit of WGS over SNP arrays is also often context-dependent, varying by trait, population, and computational methods [64]. Computational effort required for working with WGS is also a major challenge in itself.

One way forward with WGS data that is gaining traction in population and human genetics is to use Ancestral Recombination Graphs (ARGs) to efficiently represent, store, and analyse large-scale WGS datasets [65, 66, 67, 68, 69, 70]. ARGs provide a comprehensive way to represent observed variation in a sample of genomes through past branching/coalescence, mutation, and recombination events [71, 72, 68]. Recent introductions to ARGs are provided by [73, 74, 75, 76], while [68] focuses on how to encode an ARG. An ARG can be represented as a graph, where haplotypes are treated as nodes and connections between haplotypes as edges (also called branches). The haplotypes can span a small region or whole chromosome. The branches represent the transmission of DNA between the immediate ancestor and descendant haplotypes, across one or many generations. Mutations occurring along the branches change the sequence of descendant haplotypes, while recombinations change their ancestors along a haplotype. At a specific position in the genome, an ARG can be represented as a (local) DNA tree, where the root haplotype is the most recent common ancestor of all sampled haplotypes. A sequence of such local trees then forms a whole-genome ARG and a popular way to store these is with the tree sequence data format [65, 66, 68]. While there is now an active development of quantitative genetic methods to work with ARGs and phenotypes [66, 67, 69, 77, 78, 79, 80], it is still unclear how this tree-based modelling benefits the estimation of rare and drifting mutation effects.

The aim of this contribution is to study the how this tree-based model can be used to estimate the effect of mutation events and to understand how the mutations drive variation in haplotype values and downstream phenotype values. To this end, we organise this contribution in three parts. First, we study a statistical model of haplotype values on a local DNA tree with branch effects representing the sum of all mutation effects along the branch in that specific genome sequence context and hence also generation or population context. We demonstrate this model with a small example and highlight its key features. Second, we extend the demonstration with a simulation study based on the tree of mitochondrial DNA (mtDNA) from the 1000 Bull Genomes Project data [81, 82], where we consider multiple scenarios of mutation effects and study how well we can estimate them. Finally, we apply the model to a real mtDNA and phenotype dataset [83]. Throughout, we focus on a simplified case of a non-recombining region of the genome and ignore recombination, which is a focus of other work [78, 77, 79]. Results show that leveraging information from a local DNA tree can improve the estimation accuracy, provide more interpretable results, and speed up the computations.

## Methods

In this section, we develop a statistical model that works with phenotype, genetic, and haplotype values, and branch and mutation effects on a local DNA tree and demonstrate it with a small example. We then evaluate the model on simulated phenotype data based on the mtDNA tree from the 1000 Bull Genomes Project [81, 82], as well as on real phenotypic data associated with the real mtDNA from [83].

### Theory

#### Phenotype value model

Following [8], we model the observed phenotype value of an individual *y*_*i*_ as a linear combination of the population mean *µ*, genetic value *a*_*i*_ for the genome region of interest, and residual 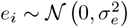 with 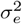 the residual variance:

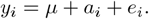

We focus on additive genetic values only and call them genetic values for brevity, but note that these are statistical parameters that depend on additive and non-additive genetic effects, and allele and genotype frequencies in a population [8, 9, 16]. For a whole data set of *n*_*y*_ phenotype records, the model in matrix form is:

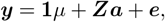

where **1** is an *n*_*y*_ *×* 1 design vector of ones for the population mean, ***Z*** is an *n*_*y*_ *× n*_*i*_ design matrix for genetic values, ***a*** is an *n*_*i*_ *×* 1 vector of genetic values, ***e*** is an *n*_*y*_ *×* 1 vector of residuals 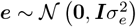, and ***a*** and ***e*** are assumed to be uncorrelated.

### Genetic value model

Let ***x***_*i*_ be an 1 *×n*_*l*_ genotype vector for individual *i* in the genome region of interest spanning *n*_*l*_ segregating biallelic loci, encoded as the dosage of alternative allele 1 compared to the reference allele 0, giving 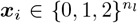. For *n*_*i*_ individuals, the corresponding *n*_*i*_ *× n*_*l*_ genotype matrix is 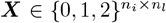. The genetic value of individual (*a*_*i*_) and the whole population of individuals (***a***) are a linear combination of individuals’ genotypes and corresponding allele substitution effects ***α*** [8, 9]:

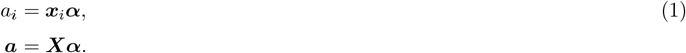

By convention, genetic values are expressed as deviations from the population mean *µ* [8], which is achieved by centring the genotypes in (1) around their expected dosage at each locus. We omit this centring from the equations for brevity and note that it does not change the estimable contrasts between model parameters.

The allele substitution effects and corresponding genetic values are estimated from observed phenotype and genotype data, commonly using the SNP-BLUP/GBLUP approach [1, 84]. To facilitate the estimation due to a large number of segregating loci and linkage-disequilibrium between them, SNP-BLUP/GBLUP assume a normal prior distribution for the allele substitution effects:

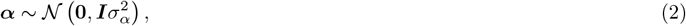

with 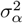 the variance of allele substitution effects. This implies a normal distribution for genetic values given the genotype matrix:

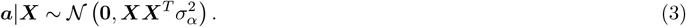

This approach assumes that allele substitution effects are constant across generations of a population and/or across populations. We aim to relax this assumption using information from a local DNA tree of haplotypes.

### Haplotype value model

The genetic value of a diploid individual for a genome region of interest is a sum of its haplotype values *h*_*i*,1_ and *h*_*i*,2_:

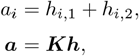

where ***K*** is an *n*_*i*_ *×* 2*n*_*i*_ design matrix between genetic values ***a*** and their underlying haplotype values ***h***. Following (1) we have for the *k* = 1, 2-th haplotype:

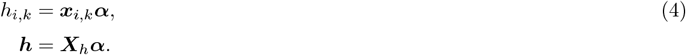

with haplotype vector 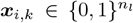 and haplotype matrix 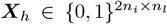. Unconditionally on the haplotype vectors, the haplotype values are normally distributed as:

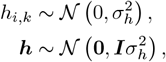

with 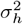 the variance of haplotype values. Conditionally on the haplotype vectors, the haplotype values are normally distributed as:

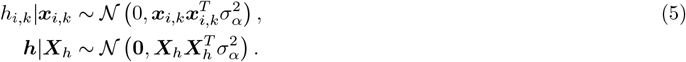

#### Haplotype value model on a local DNA tree

To expand the model (4)-(5) such that it could account for rare and drifting mutation effects across generations or populations, we consider a local DNA tree of haplotypes that are separated by mutations and corresponding changes in haplotype values due to mutation effects [85, 86]. A hypothetical example of a local DNA tree is shown in Figure 1. The aim of this expansion is driven by the fact that a local DNA tree is the generative model of haplotypes in line with the biology of DNA [71, 72]. We would like to evaluate the usefulness of this generative model also for haplotype values that underlie genetic values. To this end, we introduce four changes. First, we switch from 0*/*1 encoding of reference and alternative alleles to 0*/*1 encoding of ancestral and derived (mutation) alleles, implying that we know the ancestral allele. Second, we assume we know the local DNA tree of haplotypes, which we will use in the following. Third, we allow for multiple mutation events at the same locus, such as recurring mutations between two alleles and/or as mutations to multiple alleles, and we store these events by expanding the haplotype vector. This extended haplotype vector will have a length of *n*_*l*_ + (*n*_*m*_ *− n*_*l*_) for *n*_*l*_ loci and *n*_*m*_ mutation events, which have generated variation across the haplotypes at these loci. To avoid confusion, we use the terms *mutation haplotype vector* (***w***_*i,k*_) and *mutation haplotypes matrix* (***W*** ) with mutation event dosages, as compared to *allele haplotype vector/matrix* (***x***_*i,k*_*/****X***_*h*_) and *allele genotype vector/matrix* (***x***_*i*_*/****X***) with allele dosages. Also, for simplicity we refer to the *n*-th mutation haplotype vector as ***w***_*n*_ and to its value as *h*_*n*_. All these three extensions require additional inferences from genomic data (ancestral alleles and local DNA tree with corresponding mutation events). Fourth, we assign effects to mutations and assume they are normally and independently distributed:

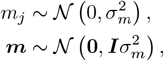

with 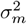 being the variance of mutation effects. This assumption is the “house of cards” model [87], where each mutation event has an independent effect of the other mutations (at the same or different loci). Other assumptions are possible [85, 88]. This setup gives a similar SNP-BLUP/GBLUP setting as in (1)-(5):

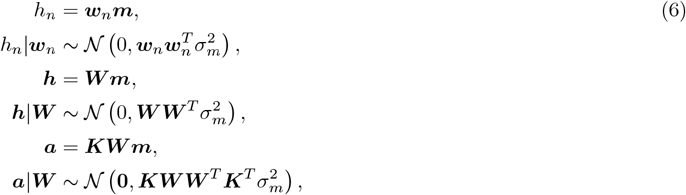

but with the mutation haplotype matrix from a local DNA tree of individuals’ haplotypes (***W*** ) instead of the matrix of “flattened” allele haplotype/genotype matrix from genotyped individuals without reference to the local DNA tree (***X***_***h***_, ***X***). We refer to this new approach as TBLUP (tree BLUP) due to leveraging the local DNA tree structure.

**Figure 1.**
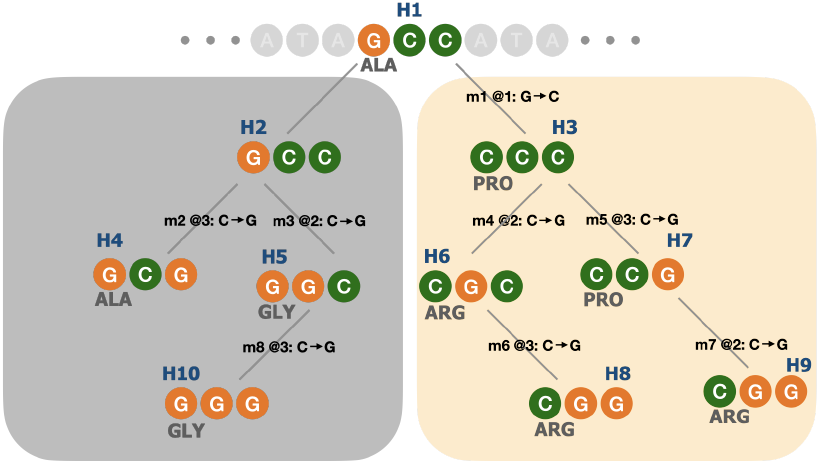
A small hypothetical local DNA tree. Haplotypes (H1-H10) span a codon in a protein-coding DNA sequence from two clades (shown as coloured boxes representing two populations) connected via the most recent common ancestor (root) haplotype (H1). Mutations between haplotypes are represented by the letter ‘m’ and sequential number, position in the codon, and nucleotide substitution; m1@1: G →C is the first mutation, it occurred at position 1, and nucleotide G mutated to C. The mutations change the codon sequence and corresponding amino acid as shown (ALA - alanine, PRO - proline, GLY - glycine, and ARG - arginine).

The mutation haplotype matrix ***W*** in TBLUP stores information on which haplotypes have inherited which mutations. Because mutations are hierarchically inherited between haplotypes, we can leverage the local DNA tree embedded in ***W*** to gain further insights and facilitate more efficient computations. Let haplotype 2 be a descendant of haplotype 1. Considering their values independently and as a function of haplotype 1 value we have:

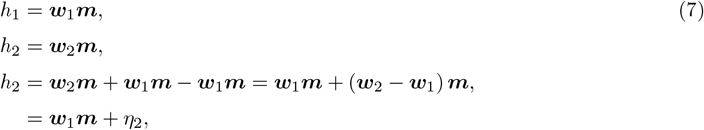

where *η*_2_ = (***w***_2_ *−* ***w***_1_) ***m*** is the change in *h*_2_ compared to *h*_1_ due to mutations that occured between haplotypes 1 and 2. In case of a single mutation, this change is simply the mutation effect *m*_*j*_, while in case of multiple mutations, this change is the sum of corresponding mutation effects. We will refer to this change in haplotype values also as the branch effect. Thence, the conditional expected value of haplotype 2 is the value of haplotype 1, while the conditional variance is proportional to the number of mutations between the two haplotypes:

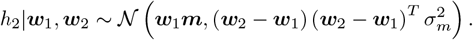

Leveraging this hierarchical local DNA tree structure and conditional independence of mutation effects also allows us to construct a sparse precision matrix for the haplotype values 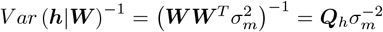 directly from the local DNA tree as shown by [85] and demonstrated in Supplement 1. In this case, we don’t centre the mutation dosages and hence the haplotype values are expressed as deviations from the root haplotype value. Working with the sparse precision matrix ***Q***_*h*_ in TBLUP is computationally advantageous over the dense precision matrix from the allele haplotype matrix 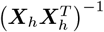 or the allele genotype matrix (***XX***^*T*^)^-1^.

We demonstrate the different approaches using allele, mutation, and local DNA tree information with the small example in Supplement 1. The demonstration is summarised with accuracy as a correlation between the true and estimated haplotype values in Table 1, showing that modelling with the mutation haplotype matrix (***W*** ) can improve the accuracy compared to modelling with the allele haplotype matrix (***X***_*h*_).

**Table 1.** Correlation between true and estimated haplotype values for the small example with different information and approaches. TBLUP uses precision matrix ***Q***_*h*_ directly from the local DNA tree, avoiding the inversion and numerical errors.

| Information | Approach | Correlation |
| --- | --- | --- |
| Allele dosage $X_h$ | SNP-BLUP | 0.87 |
|  | GBLUP | 0.87 |
| Mutation dosage $W$ | SNP-BLUP | 0.94 |
|  | GBLUP | 0.94 |
|  | TBLUP | 0.94 |

When is modelling with the mutation haplotype matrix (***W***) on a local DNA tree expected to be “helpful” compared to modelling with the allele haplotype matrix (***X***_*h*_) or allele genotype matrix (***X***)? If we consider only biallelic loci, that each allele arises from a single past mutation event, and that there are no untyped loci/variants in the genome region of interest, then the matrices ***W***, ***X***_*h*_, and ***X*** and their corresponding models work with the equivalent information. Vice versa, allele dosages in ***X***_*h*_ and ***X*** can also be polarised into ancestral and derived alleles, expanded with multiallelic loci and recurring mutations, which would provide equivalent information as ***W*** . Hence the main advantage of modelling with ***W*** over ***X***_*h*_ or ***X*** is that the underlying hierarchical generative model of haplotypes on the local DNA tree (i) provides further insights into how different mutations affect traits and (ii) facilitates faster computations with sparse matrices (***Q***_*h*_). However, critically, depending on the structure of the observed genomic and phenotypic data, we might have limited ability to estimate the mutation effects ***m*** (either with ***W*** or with ***X***_*h*_ or ***X***). To see this, note that a branch effect between haplotypes represents the sum of all mutation effects on the branch, including typed and untyped loci/variants. Hence, we can only estimate individual mutation effects if we observe haplotypes that are separated by a single mutation event and if we have associated phenotypes with these haplotypes. This is similar to estimating the Mendelian sampling term of each individual in the pedigree-based model [89]. We demonstrate this with a scenario in simulations described below.

There are two edge cases that need consideration when modelling haplotype values on a local DNA tree with TBLUP. First, the root haplotype of the local DNA tree is the most recent common ancestor of all sampled haplotypes and has by definition ancestral alleles 0 at all loci, giving a vector ***w***_0_ of zeros. This means that prior distribution (6) for the root haplotype value is “degenerate” with all the probability point mass at 0, because 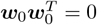. In that case the haplotype value *h*_0_ is alliased with the population mean *µ* in the model and can hence be dropped from the complete model, which then estimates other haplotype values as deviations from the root haplotype value. This is equivalent to other random walk models [90], including those used in phylogenetics [91, 92]. Second, we can observe two haplotypes in two individuals that are in ancestor-descendant relationship, but the haplotypes are identical to each other due to a lack of mutations or due to untyped loci/variants. This last case of untyped loci/variants is relevant when we work with marker genotypes from SNP arrays or reduced-representation WGS, provided that we can estimate the local DNA tree of haplotypes from the data. In this case, the descendant haplotype is an identical copy of the ancestral haplotype and its conditional prior distribution is also “degenerate” with all the probability point mass at the value of the ancestral haplotype. This is also shown as a reduced rank of the matrix ***W*** . We can address this situation in two ways. One way is to reassign any phenotype value from the descendant haplotype to the ancestral haplotype (or vice versa) and remove one haplotype from the local DNA tree and the model. Another way is to add a noise term *ϵ* with a very small variance to the descendant haplotype value (or to all haplotype values), indicating expected variation due to untyped loci/variants. This would change (7) to *h*_1_ = ***w***_1_***m*** + *ϵ*_1_ and *h*_2_ = *h*_1_ + *ϵ*_2_. The addition of the noise term also allows keeping the root haplotype in the model, because its prior variance becomes 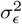.

### Application

We tested the application of the above described models and sources of information using cattle mitochondrial DNA (mtDNA). The cattle mitochondrial genome is *∼* 16 kbp long, non-recombining, maternally inherited, and haploid. Its small size allows a fast analysis and the absence of recombination means that one local DNA tree captures the genealogical history of the entire mtDNA and hence serves as a good example for demonstration. Also, being haploid, it is de-facto phased, reducing the need for phasing and limiting errors in the inference of the local DNA tree. In the following, we first describe two different mtDNA datasets, the inferrence of the ancestral alleles and the local DNA tree, simulation and empirical data analysis, finally followed by describing the evaluation of model fits from the SNP-BLUP approach on an allele haplotype matrix and the TBLUP approach on a local DNA tree. All the computations were performed on the University of Edinburgh High-Performance Computing cluster (Eddie). Scripts used in this study are available at https://github.com/HighlanderLab/gmafrafortuna_quangen_treeand use i) Python [93] for inferrence and analysis of the mtDNA tree with *tsinfer* [65, 94] and *tskit* [66, 95] packages, and ii) R [96] for data manipulation, simulation, and analysis using *Matrix* [97], *pedigreem* [98], *lubridate* [99], and *tidyverse* [100] packages. All models were fitted using the R package *INLA* [101, 102].

#### Datasets

The first dataset was derived from the publicly available 1000 Bull Genomes Project Run8 [81] (available at https://www.ebi.ac.uk/ena/browser/view/PRJEB42783). A full description of the dataset of relevance to this study is given by [82]. The dataset contained complete mtDNA sequences for 1,842 animals of *Bos taurus* and *Bos indicus* origin. The mtDNA had 4,540 sites, of which 3,518 were biallelic SNPs. The dataset had no associated phenotypes. We used the mtDNA SNPs to infer ancestral alleles and the local mtDNA tree, and to evaluate the models via simulations. Throughout the text, we refer to this dataset as *1KB*.

The second dataset was from [83]. It contained data on 359 SNPs for 96 unique mtDNA sequences of Croatian Holstein cattle, as well as the pedigree and phenotypic data for milk production traits. Due to the maternal inheritance and absence of recombination, the 96 unique mtDNA sequences linked to 3,040 females and their 7,576 lactation records through a pedigree of 6,336 animals. We used this dataset for evaluating the models via empirical data analysis. Throughout the text, we refer to this dataset as *CRO*.

#### Ancestral allele inference

To infer ancestral alleles for the cattle mtDNA, we obtained complete mtDNA sequences for seven reference species from the GenBank repository (NCBI): *Bos taurus* (V00654), *Bos indicus* (NC_005971), *Bos primigenius* (NC_013996), *Bos grunniens* (yak, KR011113), *Bison bison* (bison, NC_014044), *Ovis aries* (sheep, KR868678), and *Camelus bactrianus ferus* (camel, EF212038). We aligned the sequences with *Mafft* [103] using default settings. A poorly aligned portion of the D-loop region was discarded, giving an alignment with 16,882 bp. For alignment refinement, further exclusion of sites was performed using Gblocks 0.91b [104]. Parameters were set for the least stringent selection. The final alignment with 16,327 bp was retained for further analyses.

Cattle ancestral alleles were inferred using the software *est-sfs* [105]. Bison, yak, and camel served as the chosen outgroups due to their evolutionary relationships with cattle (the focal group). Yak and bison are closely related outgroup ruminant species, while camel, an artiodactyl, represents a more distant outgroup [106]. Allele counts for cattle were extracted from the *1KB* dataset, while the allele for each outgroup was extracted from the alignment. For each site, we obtained the probability of the major allele in the focal species being the ancestral allele. The major allele was considered ancestral if this probability was greater than 0.9, while the probability less than 0.1 indicated the minor allele as ancestral. For sites with the probability between 0.1 and 0.9 we assumed the ancestral allele to be the major allele in the outgroups. We were able to determine the ancestral state for 3,257 polymorphic sites in the mtDNA.

#### mtDNA tree inference

We inferred the mtDNA tree using the Python package *tsinfer* [65, 94]. We set the *recombination_rate* parameter to 1*e*^*−*20^ (effectively zero) and *mismatch ratio* parameter to 1*e*^18^. The mismatch ratio argument controls whether a conflict is resolved via recurrent mutation (high value, *>* 1) or recombination (low value, *<* 1). Setting the mismatch ratio to 1*e*18 forced the inference to resolve all haplotype differences through mutations instead of recombinations, as expected for the non-recombining mtDNA, resulting in a single tree with recurrent mutations, as opposed to the *tsinfer* default infinite-sites model. *tsinfer* can generate mutations above the root, especially after tree simplification that, among others, removes unary nodes [68]. To model a local DNA tree, we created a new node with all ancestral alleles and set it as an ancestor to the inferred root haplotype.

During tree sequence inference for the *1KB* dataset, we removed samples whose branches had more than 100 mutations since this could indicate, among other things, low-quality sequence data. This led to the removal of 214 samples and additional sites. The final tree had 1,684 nodes (1,669 sampled mtDNA sequences and 13 ancestors), 1,682 edges (branches), 1,029 sites, and 9,598 mutations. For the *CRO* dataset, the final mtDNA tree had 222 nodes (96 samples and 126 ancestors), 124 edges (branches), 357 sites, and 411 mutations.

#### Simulated data setup and analysis

We used the inferred mtDNA tree from the *1KB* dataset for simulation. We evaluated 36 scenarios with different assumptions about mutation effects and phenotyping scenarios. Each scenario was replicated 100 times to assess variance in results. We simulated haplotype values as the the cumulative effect of all mutations along the path from the root to a haplotype across the 1,683 nodes in the *1KB* mtDNA tree. For sample nodes (animals), we also simulated phenotype values as described above.

We considered two primary scenarios when simulating mutation effects: all mutations were causal (scenario **ALL**), or only 10% of the mutations were causal (scenario **FEW**). We then considered three sub-scenarios to determine the effect of the mutations: (i) all mutations had unique effects (**DIFF**), (ii) all mutations of the same type (such as, A *→* T) shared the same effect (**SAME**) and (iii) symmetric mutation types (such as, A *→* T and A *←* T) had the same absolute effect but opposite signs (**SYM**). In all cases, mutation effects were drawn from a normal distribution 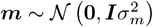 with 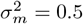. This led to six distinct mutation scenarios: ALL-DIFF, ALL-SAME, ALL-SYM, FEW-DIFF, FEW-SAME, and FEW-SYM. Having simulated mutation effects, we calculated branch effects as the sum of mutation effects along the branch between two haplotypes. Haplotype values (*h*_*n*_) were obtained by setting the root haplotype to zero *h*_0_ = 0 and recursively accumulating branch effects *h*_*n*_ = *h*_*p*(*n*)_ + *η*_*n*_, where *a*(*n*) is the immediate ancestor haplotype of haplotype *n* and *η*_*n*_ the branch effect.

We also considered three scenarios regarding phenotyping strategy: (i) all 1,669 animals were phenotyped, (ii) only a quarter of the animals (417) were phenotyped, and (iii) only 70 animals were phenotyped. Furthermore, we compared the performance of the TBLUP approach when phenotypes are available only for the 1,669 contemporary nodes (Recent), or for the contemporary and ancestral nodes (Old & Recent). However, the inferred mtDNA tree had a small number of ancestral nodes separated by many mutations on the branches connecting the nodes, representing the result of the inheritance of mtDNA across many generations. To insert the unobserved ancestral nodes, we first identified branches where mutations occurred and randomly selected one mutation per branch. We assigned a new node below each selected mutation. All mutations above the selected mutations were reassigned to the branch connecting the new node and the original parent, while mutations below the selected one were reassigned to the branch connecting the original node to the new node. This approach added 947 nodes to the initial tree, meaning that in the Recent scenario 1,669 nodes had a phenotype, while in the Old & Recent scenarion, 2,616 nodes had a phenotype. Phenotypic values were then generated just as before.

The simulated data was first modelled using the SNP-BLUP approach on column-centred *allele haplotype matrix* (***X***_*h*_) (5) to obtain estimates of marker effects (henceforth called site effects), and corresponding estimates of haplotype values. We refer to this approach as SNP-BLUP for simplicity. We next modelled the data using the TBLUP approach on *mtDNA tree* with sparse precision matrix (***Q***_*h*_) (6) to obtain estimates of haplotype (node) values, and corresponding estimates of branch effects, mutation effects, and site effects.

We evaluated the performance of the simulation and model approaches by comparing the accuracy calculated as the correlation between true and estimated effects or values. We evaluated the accuracy of haplotype values (separately for sampled haplotypes and for sampled and ancestral haplotypes), branch effects, mutation effects, and site effects. Haplotype values for ancestors, branch effects, and mutation effects were available only with the TBLUP approach. Node effects included the effect of all haplotypes (sampled and ancestral). Branch effects were estimated by multiplying the TBLUP estimates of haplotype values by ***T*** ^*−*1^ (see Supplement 1), where ***T*** is the incidence matrix connecting haplotypes (nodes) to branches. Since we could not estimate the effect of individual mutations on a branch, we naïvely estimated their effect by dividing an estimated branch effect by the number of mutations on the branch. In the opposite direction, we calculated site effects by summing the mutation effects at a locus - we did this both for the true and estimated mutation effects. We also evaluated the elapsed time for each model in both the simulation and the empirical data analysis.

#### Empirical data analysis

We refitted the SNP-BLUP model from [83] by using column-centred *allele haplotype matrix* (5) and also fitted the TBLUP approach on *mtDNA tree* (6) inferred from the *CRO* dataset. We focused on the estimation of variance components for various quantities. Both approaches used the repeatability model:

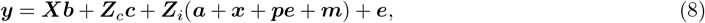

where ***y*** is the vector of milk yield records, ***X*** the design matrix for the fixed effect - age at first calving with ***b*** the corresponding regression coefficient, ***Z***_*c*_ the design matrix for the random effect of contemporary group, defined as herd-year-season effects modelled as 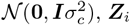 the design matrix for individual animal effects, ***a*** the vector of genetic values for the autosomal DNA modelled as 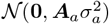 using autosomal pedigree relationship matrix ***A***_*a*_, ***x*** the vector of genetic values for the X chromosome modelled as 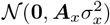 using X chromosome pedigree relationship matrix ***A***_*x*_, ***m*** the vector of genetic values for the mtDNA modelled with SNP-BLUP or TBLUP approach, ***pe*** the vector of permanent environment effects modelled as 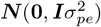, and ***e*** the vector of residuals modelled as 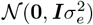.

From the fitted models (8), we obtained estimates of (“standard”) variance components listed above for SNP-BLUP and TBLUP approaches: 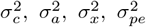, and 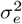. With SNP-BLUP, we also obtained a direct estimate of the variance of site effects 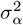. With TBLUP, we also obtained a direct estimate of the variance of mutation effects 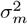. To obtain the other variance components, we used the `inla.posterior.sample` function from the *INLA* package, and obtained 1,000 Monte Carlo samples from the posterior distribution of site effects ***α*** for the SNP-BLUP approach and mutation effects ***m*** for the TBLUP approach. With these samples, we then obtained samples of haplotype values, node values, and branch effects in line with the theory and simulation subsections, and calculated their variance for each sample, giving us posterior estimates of corresponding (“derived”) variance components: 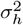 is the variance of mtDNA haplotype values (sample haplotypes), 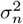 is the variance of mtDNA haplotype/node values (sample and ancestral haplotypes), and 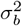 is the variance of branch effects. For each variance, we reported the posterior mean and the posterior standard deviation.

## Results

We compared the performance of the TBLUP to estimate mutation effects against the SNP-BLUP to estimate site (marker) effects in a simulation and with an empirical data analysis. In the following, we present the results for each dataset separately, showing that TBLUP tends to outperform SNP-BLUP as the number of phenotyped animals increases, but also that it reduces the computational demand, and extracts more information from the WGS data.

### Simulated data

#### Estimation of haplotype values

##### Model choice

For the majority of scenarios, both TBLUP and SNP-BLUP approaches reached comparable accuracy of estimated haplotype values Figure 2, with the average accuracy across replicates and scenarios ranging between 0.70 and 0.99. Although the differences were not large, we observed a general tendency of higher accuracies with TBLUP, particularly with more phenotyped animals. Significant differences between the approaches were observed in the scenario with 1,669 phenotypes in combination with the mutation scenarios ALL-DIFF, ALL-SYM, and FEW-DIFF. Specifically, on average, TBLUP outperformed SNP-BLUP by 6.52% in the ALL-DIFF scenario and by 9.41% in the FEW-DIFF scenario. In the ALL-SYM scenario, SNP-BLUP outperformed TBLUP on average by 4.6%.

**Figure 2.**
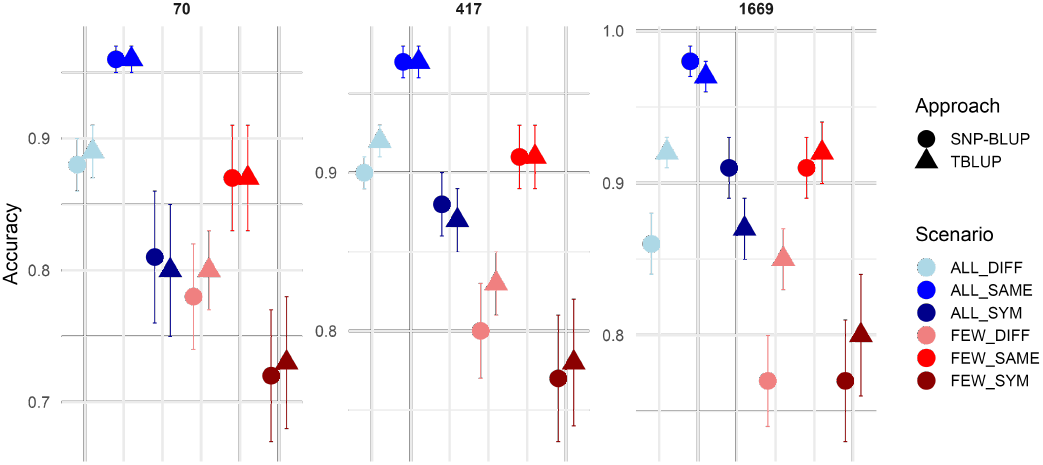
Average ( *±* standard error) accuracy of estimates for haplotype effects with SNP-BLUP and TBLUP approaches in simulation with different scenarios. The x-axis differentiates mutation scenarios, while accuracy is on the y-axis. Symbol colors and shapes distinguish mutation scenarios and the approaches: blue for all mutations having an effect (ALL), red for a few mutations having an effect (FEW), lighter colours for all mutations having different effects (DIFF), medium for the same type of mutations have the same effect (SAME), and darkest for added symmetry to SAME for reverse mutations (SYM). Circles represent SNP-BLUP and triangles TBLUP results. The three panels represent the number of phenotypes used in the analysis.

##### Phenotyping and mutation scenarios

In the majority of scenario, increasing the number of phenotyped animals improved the accuracy for TBLUP and SNP-BLUP. Increasing the number of phenotyped animals also decreased the standard error. The largest improvement was observed in the ALL-SYM scenario, where accuracy increased from the scenario with 70 animals phenptyped to all phenotyped by 0.07 (8.75%) for TBLUP and 0.1 (12.35%) for SNP-BLUP. In contrast, the ALL-DIFF and FEW-DIFF scenarios seemed to benefit less or even perform worse when the number of phenotypes increased. The ALL-SAME scenario had the highest accuracy with little difference between phenotyping scenarios. For both, TBLUP and SNP-BLUP, the accuracy ranged beween 0.70 and 0.99 with small standard error (0.01). In contrast, the FEW-SYM scenario had the lowest accuracy across all phenotyping scenarios, between 0.72 and 0.80. Overall, within each mutation scenario (ALL or FEW), scenarios with the same effects for a mutation type (ALL-SAME and FEW-SAME) consistently performed better, while scenarios with symmetric mutation effects (ALL-SYM and FEW-SYM) proved more challenging and for majority of these TBLUP performed better.

##### Elapsed time

We evaluated the elapsed time to run TBLUP and SNP-BLUP across the scenarios (Figure 3 and Table 7). Overall, the TBLUP approach was fitted faster than SNP-BLUP with *INLA*. TBLUP elapsed time was less than 3 seconds for all scenarios, while SNP-BLUP elapsed time ranged from 7.5 seconds with 70 phenotyped animals to more than 400 seconds with 1,669 phenotyped animals. There was some variation in elapsed time across mutation scenarios for SNP-BLUP with 1,669 phenotyped animals; with large average elapsed time and its standard error, as well as indication of a larger elapsed time when a few mutations were causal.

**Figure 3.**
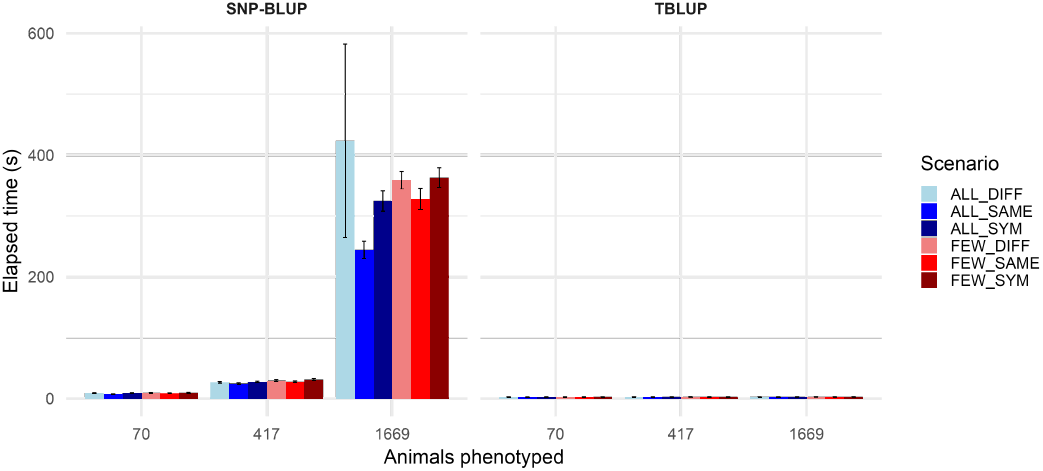
Average (*±* standard error) elapsed time in seconds for TBLUP and SNP-BLUP approaches across scenarios. The x-axis differentiates number of phentyped animals, while elapsed time is on the y-axis. Colors distinguish mutation scenarios: blue for all mutations having an effect (ALL), red for a few mutations having an effect (FEW), lighter colours for all mutations having different effects (DIFF), medium for the same type of mutations have the same effect (SAME), and darkest for added symmetry to SAME for reverse mutations (SYM).

#### Estimation of node values and branch, mutation, and site effects

In the simulation, we also assessed the accuracy of estimating node values (haplotype values for sampled and ancestral haplotypes) and decomposing these into branch, mutation, and site effects with TBLUP (Figure 4). In general, the accuracy of estimates decreased as we decomposed the node values into branch effects and then into mutation effects, while site effects had intermediate accuracy between the branch and mutation effects. Node, branch, and mutation estimates were available only with the TBLUP approach, so comparison with SNP-BLUP was only possible for the site effects.

**Figure 4.**
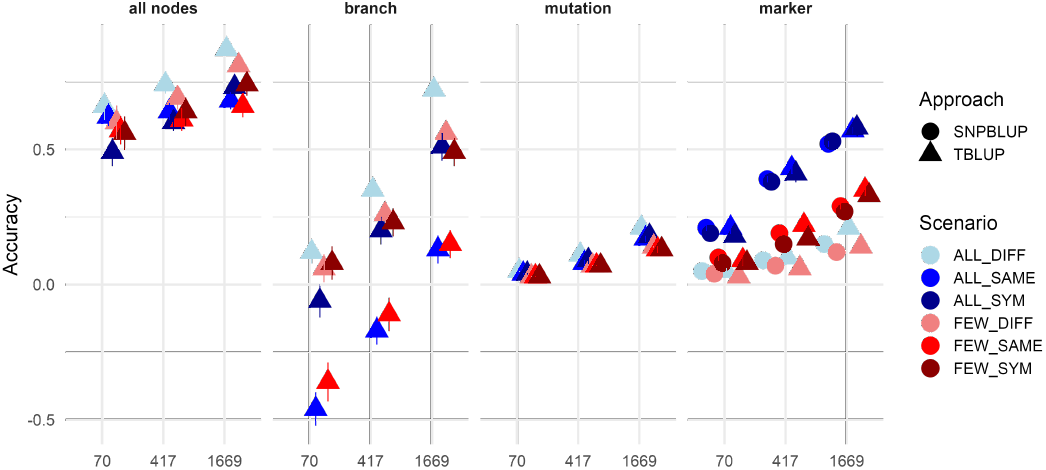
Average (*±* standard error) accuracy of estimates for node, branch, mutation, and site effects. The x-axis differentiates sample sizes, while accuracy is on the y-axis. Symbol colors and shapes distinguish mutation scenarios and the approaches: blue for all mutations having an effect (ALL), red for a few mutations having an effect (FEW), medium for the same type of mutations have the same effect (SAME), and darkest for added symmetry to SAME for reverse mutations (SYM). Circles represent SNP-BLUP and triangles TBLUP results.

##### Node effects

The accuracy of node value estimates ranged from 0.49 *±* 0.05 (for the FEW-SYM scenario with 70 animals) to 0.87*±*0.02 (for the ALL-DIFF scenario with 1,669 animals). The accuracy generally increased with more phenotyped animals, however, the rate of increase differed between mutation scenarios Figure 2. Specifically, when we increased the number of phenotyped animals from 70 to 1,669, the accuracy increased most in the ALL-SYM scenario (49.0%), followed by the increase in the FEW-DIFF (35.0%), FEW-SIM (32.1%), and ALL-DIFF scenario (31.8%).

##### Branch effects

The accuracy of estimated branch effects was the most sensitive to phenotyping scenarios and benefited the most from increasing the number of phenotyped animals across all mutation scenarios. The greatest improvement was observed when increasing the number of phenotyped animals from 70 (0.12*±*0.04) to 1,669 phenotypes (0.72 *±* 0.02) in the ALL-DIFF scenario. The ALL-DIFF scenario also had the highest accuracy for branch effects, followed by FEW-DIFF, FEW-SYM, and ALL-SYM scenarios, while the ALL-SAME and FEW-SAME had the lowest accuracy with negative accuracies with small numbers of phenotyped animals.

##### Mutation effects

The accuracy of estimated mutation effects was the least impacted by the extent of phenotyping extent. This is expected since we are not able to discriminate the effect of individual mutations on a branch and we have to assume a naïve estimate of dividing the branch effect by the number of mutations. The highest accuracy was obtained for the ALL-DIFF scenario with 1,669 phenotypes (0.21 *±* 0), which is in line with the high accuracy of estimating branch effects in this scenario.

##### Site effects

In addition to comparing the accuracy of estimated haplotype values in Figure 2, we compared TBLUP and SNP-BLUP based accuracies of estimated site effects. In general, there was no significant difference between the two approaches, with slight tendency of TBLUP to have higher accuracy estimates. Increasing the number of phenotyped animals improved the accuracy of estimating site effects for both TBLUP and SNP-BLUP; with the highest accuracy of 0.58 *±* 0.03 for ALL-SYM scenario with 1,669 phenotypes. The accuracy increased the most from the scenario with 70 to 1,669 phenotypes for the ALL-SAME and ALL-SYM scenarios, followed by FEW-SAME and FEW-SYM, with the lowest increases for ALL-DIFF and FEW-DIFF. There was no significant difference between the two approaches, with slight tendency of TBLUP to have higher accuracy estimates.

### Recent vs. Old & Recent phenotyping scenarios

We evaluated the performance of the TBLUP approach in a scenario in which phenotypes were available only for contemporary nodes (Recent) and another in which phenotypes were available also for ancestral nodes (Old & Recent). The accuracy of estimates for phenotyped haplotypes, all haplotypes, branch effects, and mutation effects for these two scenarios are presented in Figure 5. No significant differences were observed for the accuracy of estimates for phenotyped haplotypes between the phenotyping scenarios. For the mutation scenarios ALL-DIFF, ALL-SAME, ALL-SYM, FEW-DIFF, and FEW-SAME, the accuracies were close to 1. For the FEW-SYM scenario, the accuracy was 0.86. Scenario with Old & Recent phenotypes had higher accuracy then only Recent for all haplotype values, especially for the mutation scenarios ALL-SAME (0.79 *±* 0.000 vs. 0.67 *±* 0.000 in the Recent scenario), ALL-SYM (0.87 *±* 0.000 vs. 0.79 *±* 0.003 in the Recent scenario), FEW-SAME (0.80 *±* 0.000 vs. 0.70 *±* 0.000 in the Recent scenario), and FEW-SYM (0.81 *±* 0.005 vs. 0.77 *±* 0.007 in the Recent scenario). The same trend was observed for branch effects, with the Old & Recent phenotyping scenario yielding higher accuracies for all mutation scenarios. Notably, negative accuracies were obtained in the Recent phenotyping scenario in the mutation scenarios ALL-SAME and FEW-SAME (the same was observed in the previous results for branch effect estimations). With the spread of phenotypes across the tree (Old & Recent scenario), this artefact of negative accuracies disappeared and all estimates had positive accuracies. The accuracy of mutation effects improved significantly with the Old & Recent phenotyping scenario in all mutation scenarios. This increase in accuracy ranged from 0.03 in the FEW-SYM mutation scenario to 0.06 in the FEW-DIFF scenario. Similarly as observed in the previous results, the accuracy of estimated haplotype values for phenotyped nodes was the highest. Accuracies decreased progressively, from haplotype values for all nodes to mutation effects, as the number of variables to be estimated increased.

**Figure 5.**
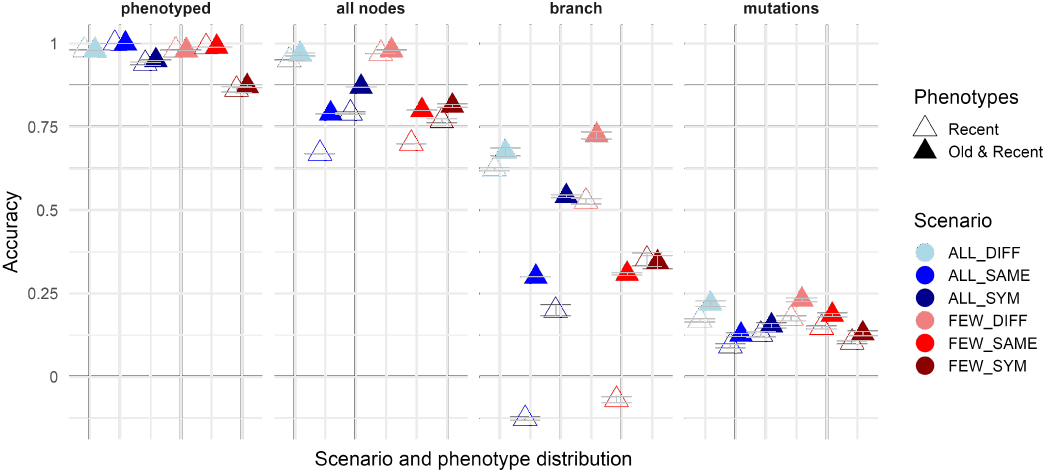
Average (*±* standard error) accuracy of estimates for phenotyped nodes, all nodes, branch, and mutation effects. The x-axis differentiates scenarios and phenotype distribution; while accuracy is on the y-axis. Symbol colors and shapes distinguish mutation and phenotyping scenario: blue for all mutations having an effect (ALL), red for a few mutations having an effect (FEW), medium for the same type of mutations have the same effect (SAME), and darkest for added symmetry to SAME for reverse mutations (SYM). Hollow triangles represent the scenario with phenotypes only for recent nodes, while solid triangles represent the scenario witj phenotypes for both old and recent nodes.

### Empirical analysis

We compared the TBLUP and SNP-BLUP approaches for the estimation of variance components using the real *CRO* dataset (Table 2). Both approaches estimated similar variance components with some minor differences. SNP-BLUP approach estimated the genetic variance for autosomal DNA 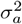 at 0.324 *±* 0.010, while TBLUP estimated it at a larger 0.377 *±* 0.010. Correspondingly, SNP-BLUP approach estimated the residual variance 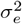 at 0.376 *±* 0.028, while the TBLUP estimate was lower at 0.330 *±* 0.027. Variance of site effects 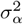 was estimated at 0.002 *±* 0.005 with SNP-BLUP, while the variance of mutation effects 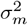 was estimated to be larger at 0.004 *±* 0.001 with TBLUP. The variance of haplotype values 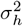 was estimated at 0.074 *±* 0.013 with SNP-BLUP, and at a slightly lower 0.065 *±* 0.012 with TBLUP, which equaled the estimate of the variance of all haplotype (node) values 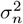 with TBLUP. Variance of branch effects 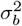 was estimated at 0.063 *±* 0.012 with TBLUP. For our data (359 SNPs for 3,040 animals) and this model complexity, the elapsed time to fit the TBLUP was on average 74.7 seconds, while for SNP-BLUP it was 77.9 seconds.

**Table 2.** Estimated variance components from the *CRO dataset*. The “standard” components are: 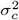 is the contemporary group (herd-year-season) variance, 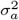 is the genetic variance for autosomal DNA, 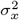 is the genetic variance for X chromosome,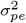 is the permanent environment variance, 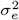 is the residual variance. The SNP-BLUP and TBLUP specific effect components are: 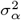 is variance of site effects and 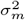 is the variance of mutation effects. The “derived” components are: 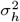 is the variance of mtDNA haplotype values (sample haplotypes), 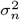 is the variance of mtDNA haplotype/node values (sample and ancestral haplotypes), and 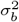 is the variance of branch effects.

| Variance components | SNP-BLUP | TBLUP |
| --- | --- | --- |
| “Standard” |  |  |
| $\sigma_c^2$ | 0.096 ± 0.008 | 0.095 ± 0.009 |
| $\sigma_a^2$ | 0.324 ± 0.010 | 0.377 ± 0.010 |
| $\sigma_x^2$ | 0.000 ± 0.000 | 0.000 ± 0.000 |
| $\sigma_{pe}^2$ | 0.061 ± 0.018 | 0.057 ± 0.017 |
| $\sigma_e^2$ | 0.376 ± 0.028 | 0.330 ± 0.027 |
| SNP-BLUP and TBLUP specific |  |  |
| $\sigma_\alpha^2$ | 0.002 ± 0.005 | NA |
| $\sigma_m^2$ | NA | 0.004 ± 0.001 |
| “Derived” |  |  |
| $\sigma_h^2$ | 0.074 ± 0.013 | 0.065 ± 0.012 |
| $\sigma_n^2$ | NA | 0.065 ± 0.011 |
| $\sigma_b^2$ | NA | 0.063 ± 0.012 |

## Discussion

The results show that, in a non-recombining DNA region, the presented TBLUP approach tends to outperform SNP-BLUP in terms of computational effort, estimation accuracy, and informativeness of results. Computational efficiency comes from the generative model of haplotype values on a local DNA tree and associated conditional distributions for ancestor–descendant haplotype values with a sparse precision matrix. Generally, estimation accuracy tended to be higher with the TBLUP approach, however, this benefit depends on the additional information of mutation dosages compared to allele dosages, and on the data structure with respect to phenotype distribution across the sampled and ancestral haplotypes in the local DNA tree. Using a local DNA tree with associated branches and mutations, the TBLUP approach also provides additional information about branch effects and mutation effects, shedding light on the estimability of these quantities and how they drive variation in haplotype and phenotype values. In the following, we discuss three points that arise from this work: (i) novel contributions and the advantages of the TBLUP approach; (ii) why are mutation effects difficult to estimate; (iii) opportunities and future directions with local DNA trees and ARGs.

### Novel contributions and the advantages of TBLUP

The novelty of the TBLUP approach lies in its ability to leverage the topology of a local DNA tree and the detailed ancestral information encoded within it. This provides a rich framework for modelling genetic variation in a non-recombining genomic region, leading to three main advantages. First, the estimation accuracy can exceed the SNP-BLUP approach, especially when phenotypes are available for recent and ancestral haplotypes. When only recent haplotypes are phenotyped, the TBLUP accuracies are comparable to the SNP-BLUP because there is no information to estimate the effect of past mutations from recent haplotypes and their phenotypes in a region of non-recombining DNA. When recombination is present, using TBLUP allows us to better localise all causal loci in the haplotype, though we still can not estimate the effect of past mutations. Research on the modelling with recombining DNA is addressed elsewhere [77, 78, 79, 80] When phenotypes are available for recent and ancestral haplotypes, the TBLUP approach achieves significantly higher accuracies. It does so because it can estimate the effect of past mutations that differentiate ancestral and descendant haplotypes and hence estimate their effect at the point they occur, though the estimates are shared across the local DNA tree. Although we can not gather phenotypes for past individuals, there is an opportunity that routine phenotyping and genotyping will enable the estimation of *de-novo* mutations going forward. Second, the TBLUP approach is based on a richer model. It provides estimates for the effect of sampled haplotypes, as well as for the ancestral haplotypes, branches between all the haplotypes, and associated mutations. These estimates provide a more detailed view of the genetic architecture underlying trait variation and how it evolved over time and is hence akin to quantitative genomic models that aim to capture recent mutations in addition to past mutations [107, 108]. Also, the branch effect variance is proportional to the mutational variance [109, 79]. Lastly, the TBLUP approach offers computational advantages. By incorporating the structure of a local DNA tree into the BLUP framework, we replace the dense genomic relationship matrix between individuals/haplotypes with a sparse graph representation of DNA inheritance between generations. As genomic datasets grow, computing, storing, and inverting genomic relationship matrices is increasingly limiting, though various approaches have been proposed to overcome this problem [110, 111, 112, 113, 114]. The TBLUP approach bypasses the need to setup and invert the relationship matrix, by directly constructing the sparse inverse (precision), **Q**_*h*_, akin to the pedigree-based model [115]. We have previously worked on this problem [85] as have others in the context of phylogeny [116, 91]. The phylogenetic models work with the whole genome and are used to model variation between (sub-)populations, while our work is for a region of DNA and models variation between individuals (within or across populations). Here, we have now evaluated the benefits of the generative modelling approach more extensively, showing the potential and possibility of estimating context-dependent effects (see also [86]). This work is part of a larger effort to leverage ARGs in quantitative genetics [66, 67, 69, 77, 78, 79], which rests on the efficiency of computing with large-scale genomic datasets stored in the tree sequence data format [65, 66, 68].

### Why are mutation effects difficult to estimate

One of the hypotheses of this study was that the TBLUP approach could accurately estimate mutation effects. Our results show that this remains challenging, even when ancestral haplotypes are phenotyped. Two main factors justifying this limitation are: (i) small size of our dataset (although we focused on just one local DNA tree) and (ii) complete linkage inherent to a non-recombining genome region. Under these conditions, mutations share similar inheritance patterns, which diversify over generations, but slowly. This condition makes it difficult to distinguish mutation effects, particularly if a group of mutations is observed on the same branch of a local DNA tree. In our study, we simply approximated mutation effects by dividing the estimated branch effect by the number of mutations observed on it. Furthermore, we had more mutations than phenotypes (9,598 vs 1,669), so our statistical power was limited. These results echo the warnings about the interpretation of past genomic trends [117]. They also highlight a fundamental constraint of working with non-recombining genome regions and motivate future work with recombining genome regions. Since recombinations reduce the linkage-disequlibrium between sites and hence mutations, we expect to be able to better disentangle the effects of mutations in recombining regions.

### Opportunities and future directions with local DNA trees and ARGs

Despite the advantages and novelty of this study, several challenges and opportunities remain. The TBLUP approach requires an accurate reconstruction of a local DNA tree, and ultimately complete ARGs to work across the whole genome. Errors in this reconstruction may propagate into the statistical estimation of the effects. The field of ARG inference is advancing rapidly, with many scalable algorithms now available [70, 118, 94, 65, 119, 120]. As these methods improve, so will the applicability of TBLUP and similar approaches. Although the sparse precision matrix provides computational benefits, the magnitude of these gains depends on software implementation. Here we have used R-INLA [101, 102] to fit the models because it is optimised for work with sparse matrices, but other software optimised for dense matrices may perform better. Nevertheless, setting up a sparse precision matrix remains an advantage compared to calculating and inverting a dense genomic relationship matrix. The accuracy of estimated mutation effects will improve by working with the full ARG that encompases all local DNA trees and hence recombinations between these. There is an active research about this topic [67, 69, 77, 78, 79] While we focused on a non-recombining region of genome, this study demonstrates the potential and benefits of leveraging the generative model for haplotype effects in quantitative genetics.

## Conclusions

In summary, the TBLUP approach that leverages information from a local DNA tree with associated mutation events provides a promising extension of the current SNP-BLUP and GBLUP approaches in terms of computational speed, estimation accuracy, and information content (estimation of haplotype values, branch effects, and mutation effects). As such, it estimates the effects of mutations in the context of a local DNA tree and enables their estimation at the point of occurrence, with the possibility to capture non-additive effects if the same type of mutation occurs in different parts of the tree. Although we studied the TBLUP approach in a non-recombining region of the genome, its potential is even greater when applied across recombining regions of the genome, which is the focus of other research [77, 78, 79, 80]. Further extension of the approach and its evaluation in more challenging settings will be crucial to fully understanding its capabilities and limitations.

## Acknowledgements

Not applicable.

## Availability of data and materials

All scripts for ARG inference, simulations, and real data analysis are available at https://github.com/HighlanderLab/gmafrafortuna_quangen_tree.

## Ethics approval and consent to participate

Not applicable.

## Consent for publication

Not applicable.

## Authors’ contributions

GMF co-designed the study, performed the analyses, interpreted results, and drafted the manuscript; JO and AM contributed to the analysis of 1000 Bull Genomes mtDNA and manuscript revision; and GG initiated and supervised the study, co-designed the study, developed theory, contributed to interpretation and writing. All authors read and edited manuscript drafts, and approved the final manuscript.

## Additional Files

Additional file 1 — Small example with demonstration of the modelling approaches. Small example and demonstration of the modelling approaches

In this section, we demonstrate the model of haplotype values on a local DNA tree with a small example. This example illustrates how mutation effects can be estimated in their genome sequence context when observed alleles are polarised into ancestral and derived states (mutations), and the local DNA tree with corresponding mutations between haplotypes is known. We fit the standard SNP-BLUP and GBLUP approaches with the allele haplotype matrix and with the mutation haplotype matrix, and the TBLUP approach with the local DNA tree. We consider two phenotyping scenarios: (i) contemporary and ancestral haplotypes have phenotypes and (ii) only contemporary haplotypes are phenotyped. R code for the calculations is available at https://github.com/HighlanderLab/gmafrafortuna_quangen_tree.

Consider a hypothetical local DNA tree in Figure 1, which describes the relationship among nine haplotypes across two clades (each denoted with a coloured box representing two populations) and their most recent common ancestor (root) haplotype. These haplotypes span one codon in a protein-coding region of DNA, hence each haplotype is represented by three nucleotides that encode an amino acid. The root haplotype (H1) has the codon GCC, encoding alanine (ALA). The tree shows a hypothetical evolutionary history of the haplotypes over a long time period, where mutations changed the codon sequence and the corresponding amino acid. For example, H3 inherited from H1 with GCC codon (alanine), but mutation 1 at site 1 has changed the codon to CCC (proline). In total, we have ten haplotypes separated by eight mutations, and these haplotypes encode four different amino acids listed in (Table 3): alanine (ALA), proline (PRO), glycine (GLY), and arginine (ARG). The example is deliberately extreme with respect to the amount and type of mutations, to emphasise the potential for modelling. The 10 *×* 3 allele haplotype matrix ***X***_*h*_ (columns A1, A2, and A3 in Table 3) and the 10 *×* 8 mutation haplotype matrix ***W*** for this example are:

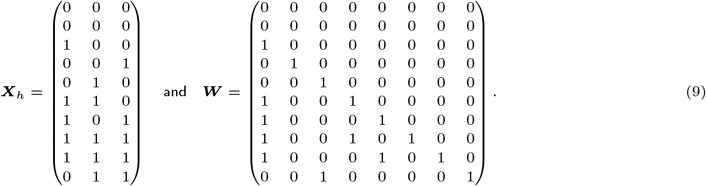

**Table 3.** Haplotype information for the small example. Information on the ten haplotypes in the local DNA tree (Haplotype) from Figure 1, their immediate ancestor haplotype (Ancestor), ancestral/derived allele encoding at the three nucleotides (A1, A2, and A3), amino acid (Amino Acid: ALA - alanine, PRO - proline, GLY - glycine, ARG - arginine), occurrence of a mutation since the ancestor (Mutated), mutated site (Site), mutation (Mutation), mutation effect (Effect), and haplotype value (Value).

| Haplotype | Ancestor | A1 | A2 | A3 | Amino acid | Mutated | Site | Mutation | Effect | Value |
| --- | --- | --- | --- | --- | --- | --- | --- | --- | --- | --- |
| H1 | / | 0 | 0 | 0 | ALA | 0 | / | / | / | 0 |
| H2 | H1 | 0 | 0 | 0 | ALA | 0 | / | / | / | 0 |
| H3 | H1 | 1 | 0 | 0 | PRO | 1 | 1 | m1 | 1 | 1 |
| H4 | H2 | 0 | 0 | 1 | ALA | 1 | 3 | m2 | 0 | 0 |
| H5 | H2 | 0 | 1 | 0 | GLY | 1 | 2 | m3 | 3 | 3 |
| H6 | H3 | 1 | 1 | 0 | ARG | 1 | 2 | m4 | 1 | 2 |
| H7 | H3 | 1 | 0 | 1 | PRO | 1 | 3 | m5 | 0 | 1 |
| H8 | H6 | 1 | 1 | 1 | ARG | 1 | 3 | m6 | 0 | 2 |
| H9 | H7 | 1 | 1 | 1 | ARG | 1 | 2 | m7 | 1 | 2 |
| H10 | H5 | 0 | 1 | 1 | GLY | 1 | 3 | m8 | 0 | 3 |

For example, H3 has inherited mutation 1, H6 has inherited mutations 1 and 4, and H8 has inherited mutations 1, 4, and 6 (Figure 1). The relationship between ***X***_*h*_ and ***W*** in this example is such that adding up columns of ***W*** for mutations at the same site gives the corresponding site columns of ***X***_*h*_. For example, there is one mutation at site 1 (mutation 1), so the first columns of ***X***_*h*_ and ***W*** are the same. There are three mutations at site 2 (mutations 3, 4, and 7), so adding these columns of ***W*** gives the second column of ***X***_*h*_. In the case of reverse mutations, this relationship does not hold for the encoding used. Coding reverse mutations in ***W*** as −1 would allow the relationship to hold. As noted in the main text, it is common to centre the columns of ***X***_*h*_ and ***W*** around the expected dosage at each locus [1, 84]. We omit this centring for brevity and note that it does not change the estimable contrasts between model parameters. It will also highlight the potential of working with the sparse precision matrix with the TBLUP approach for haplotype values.

We assume that each amino acid has an effect on a trait of interest and therefore each underlying haplotype has a value for the trait. We assume that the root haplotype (H1) has a value of 0 and will hence be aliased with the model intercept. The derived haplotypes are assigned different values compared to the root in line with their amino acid. We arbitrarily assigned these haplotype values to the amino acids: alanine (ALA = 0, root), glycine (GLY = 3), arginine (ARG = 2), and proline (PRO = 1), which also induces corresponding mutation effects (Table 3). Note that these mutation effects depend on the genome context (surrounding nucleotides) and hence manifest non-additive effects. For example, mutation 3 at site 2 (C →G) changed alanine to glycine with the corresponding effect of 3 units, while mutations 4 and 7 at site 2 (also C →G) changed proline to arginine with the corresponding effect of 1 unit. Allele substitution effects ***α*** at the three nucleotides calculated independently (by fitting ***h*** = **1***µ* + ***X***_*h,k*_*α*_*k*_ + ***e*** for each column *k* of ***X***_*h*_) are 0.4, 2.0, and 0.4, while calculated jointly (by fitting ***h*** = **1***µ* + ***X***_*h*_***α*** + ***e***) are ∼ 0.0, 2.0, and ∼ 0.0.

Based on this local DNA tree, we generated a small dataset by randomly sampling the haplotypes for nine individuals and simulating phenotypes for the individuals from ***y*** = **1***µ* + ***Zh*** + ***e***, where *µ* = 10, ***h*** are haplotype values from Table 3, and ***e*** ∼ *N* (0, 1). The simulated data are shown in Table 4.

**Table 4.**
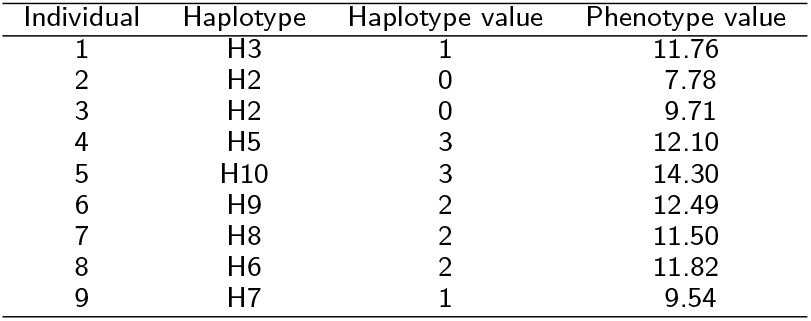
Simulated data based on the small example for the first phenotyping scenario. In this scenario ancestral and contemporary haplotypes have associated phenotypes.

| Individual | Haplotype | Haplotype value | Phenotype value |
| --- | --- | --- | --- |
| 1 | H3 | 1 | 11.76 |
| 2 | H2 | 0 | 7.78 |
| 3 | H2 | 0 | 9.71 |
| 4 | H5 | 3 | 12.10 |
| 5 | H10 | 3 | 14.30 |
| 6 | H9 | 2 | 12.49 |
| 7 | H8 | 2 | 11.50 |
| 8 | H6 | 2 | 11.82 |
| 9 | H7 | 1 | 9.54 |

In the following, we demonstrate the key steps with SNP-BLUP and GBLUP approaches with allele haplotype matrix ***X***_*h*_ and mutation haplotype matrix ***W*** (the TBLUP approach).

### SNP-BLUP with allele dosages and phenotypes for all haplotypes

Using the *SNP-BLUP approach* (2) with the phenotype values ***y*** and the design matrix ***Z*** from Table 4, the *allele haplotype matrix* ***X***_*h*_ from (9), and assuming 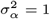 and 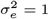, we get the following SNP-BLUP system of equations and estimates:

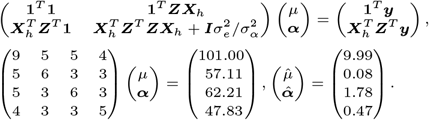

The estimates of allele substitution effects 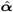 differ from the true values above due to the small noisy dataset and penalised estimation. Applying these estimates to the allele haplotype matrix, gives estimates of haplotype values:

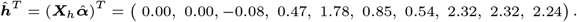

These estimates of haplotype values differ from the true values in Table 3 because we used only a small noisy dataset in the estimation (Table 4), but also because the SNP-BLUP approach uses multiple linear regression on allele dosages, while the mutation effects in the local DNA tree have additive and non-additive effects.

### GBLUP with allele dosages and phenotypes for all haplotypes

Using the *GBLUP approach* (5) with the phenotype values ***y*** and the design matrix ***Z*** from Table 4, the *allele haplotype matrix* ***X***_*h*_ from (9), and assuming 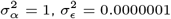, and 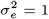, we get the following GBLUP system of equations and estimates:

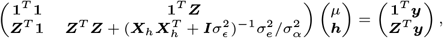

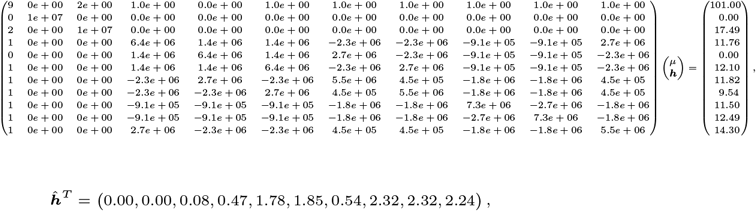

which are the same estimates of haplotype values as with the SNP-BLUP approach with allele dosages.

### SNP-BLUP with mutation dosages and phenotypes for all haplotypes

Using the *SNP-BLUP approach* (2) with the phenotype values ***y*** and the design matrix ***Z*** from Table 4, the *mutation haplotype matrix* ***W*** from (9), and assuming 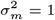 and 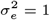, we get the following SNP-BLUP system of equations and estimates:

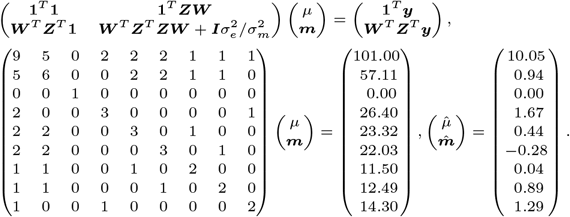

The estimates of mutation effects 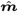 resemble the true effects, but ultimately differ due to the small noisy dataset. Applying these estimates to the mutation haplotype matrix, gives estimates of haplotype values:

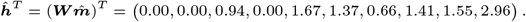

### GBLUP with mutation dosages and phenotypes for all haplotypes

Using the *GBLUP approach* (3) with the phenotype values ***y*** and the design matrix ***Z*** from Table 4, the *mutation haplotype matrix* ***W*** from (9), and assuming 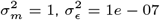, and 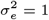, we get the following GBLUP system of equations and estimates:

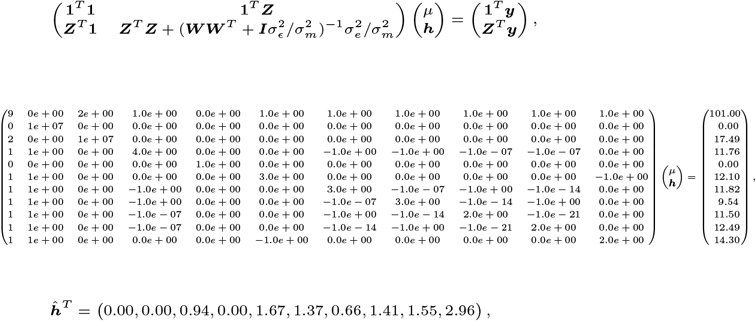

which are the same estimates of haplotype values as with the SNP-BLUP approach with mutation dosages. Critically, the left-hand-side of the above system of equations is sparse, because the precision matrix for haplotype values is sparse; 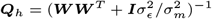, eventhough the corresponding covariance matrix 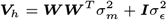 is dense.

### TBLUP with a local DNA tree and phenotypes for all haplotypes

Instead of solving for the ***Q***_*h*_ matrix, incurring a computational cost and numerical errors, we can set it directly from the local DNA tree [85] similar to the pedigree model [89]. This follows from the hierarchical generative model (6):

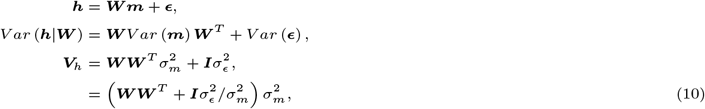

Generalized Cholesky decomposition of the covariance matrix ***V***_*h*_ and its inverse is then:

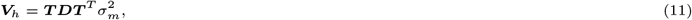

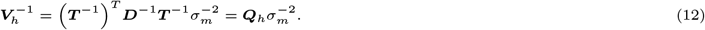

For the small example the triangular matrix ***T*** ^−1^ is:

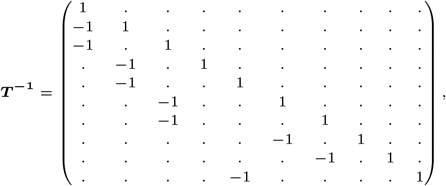

where diagonal elements are 1, non-zero off-diagonal elements are −1 for each pair of immediate descendant-ancestor haplotypes, and all other elements are 0, hence directly reflecting the local DNA tree. For example, H2 and H3 are immediate descendants of H1, while H4 and H5 are immediate descendants of H2, etc. The diagonal matrix ***D*** is conditional variance of haplotype values, given the ancestor haplotype value, hence the variance of the branch effects (the sum of mutation effects on each branch) plus variance of additional terms added to haplotype values (*ϵ*). For the root haplotype, this is equal to:

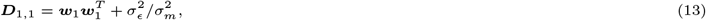

while for any other haplotype *i* it is,

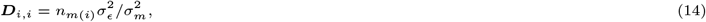

where *n*_*m*(*i*)_ is the number of mutations that separate the haplotype from its immediate ancestor. With a way to obtain ***T*** ^−1^ and ***D***^−1^ efficiently directly from the local DNA tree, we can also efficiently setup ***Q***_*h*_. For the small example, this precision matrix is:

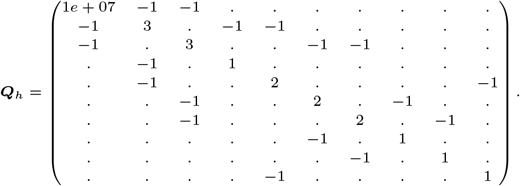

Using the above precision matrix ***Q***_*h*_ instead of inverting the covariance matrix ***V***_*h*_, produced these estimates of haplotype values:

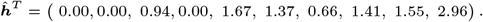

To obtain estimates of mutation effects from the estimates of haplotype values, we can pre-multiply the latter with 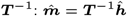

### Comparison of the estimates across the approaches and phenotyping scenarios

In addition to Table 4 with phenotypes for all haploptypes, we have also fit GBLUP with an allele dosage matrix and TBLUP with the local DNA tree with phenotypes for contemporary haplotypes only shown in Table 5. To facilitate the comparison between the two phenotyping scenarios, the number of phenotypes was kept the same for both scenarios. In Table 6 we compare true haplotype values with their estimates from GBLUP with an allele dosage matrix and TBLUP with the local DNA tree, true mutation effects with their estimates from TBLUP, and true site effects with their estimates from the GBLUP are also provided and compared between phenotyping scenarios. The table shows that the accuracy of predictions increases using the TBLUP approach when ancestral haplotypes are also phenotyped (from 0.81 with only contemporary haplotypes phenotyped to 0.94 when ancestral haplotypes are phenotyped). The power of the TBLUP approach to estimate the effect of mutation decreases when ancestral haplotypes are not phenotyped, with accuracy reducing from 0.75 (when ancestral and contemporary haplotypes are phenotyped) to 0.45 (when only contemporary haplotypes are phenotyped).

**Table 5.** Simulated data based on the small example for the second phenotyping scenario. In this scenario only contemporary haplotypes (at the tips of the tree) have associated phenotypes.

| Individual | Haplotype | Haplotype value | Phenotype value |
| --- | --- | --- | --- |
| 1 | H8 | 2 | 13.18 |
| 2 | H8 | 2 | 11.41 |
| 3 | H8 | 2 | 13.46 |
| 4 | H9 | 2 | 13.69 |
| 5 | H4 | 0 | 11.22 |
| 6 | H10 | 3 | 13.33 |
| 7 | H9 | 2 | 10.87 |
| 8 | H8 | 2 | 13.10 |
| 9 | H8 | 2 | 12.94 |

**Table 6.**
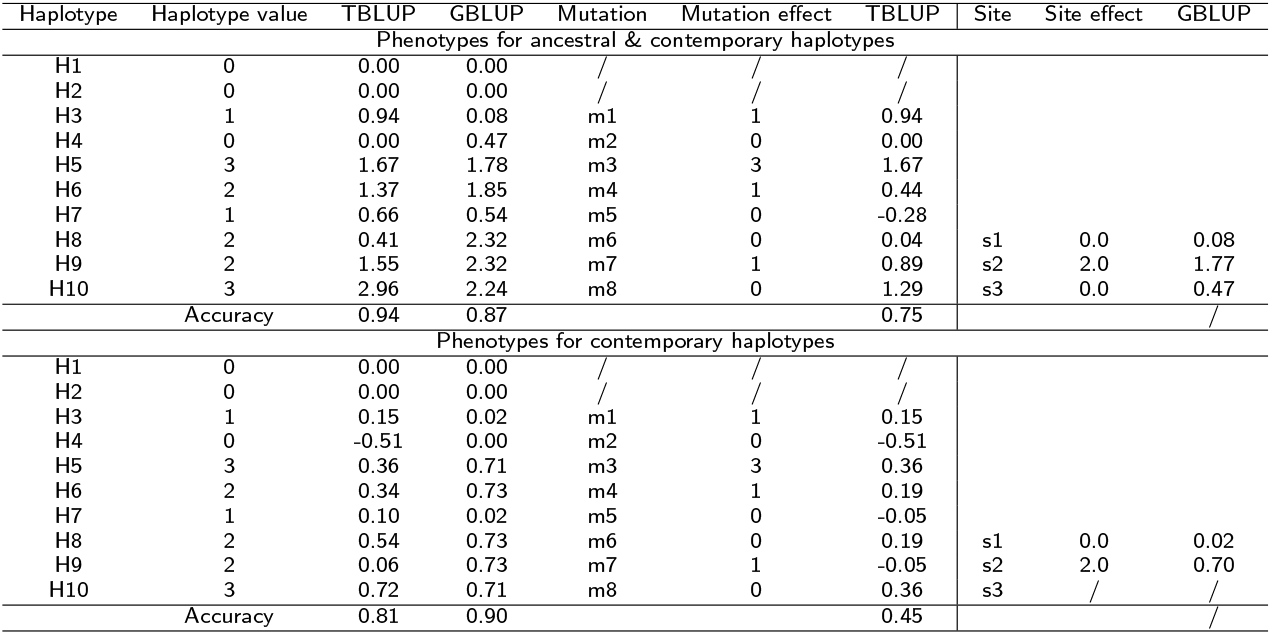
Comparison of the true and estimated haplotype values and mutation effects across approaches and phenotyping scenarios. Estimates were obtained from GBLUP approach on allele dosage matrix and TBLUP approach on the local DNA tree with two phenotyping scenarios. Site effects are obtained from a joint regression of estimated haplotype effects on the allele dosage matrix. In the second phenotyping scenario (phenotypes for contemporary haplotypes only) allele effects can be estimated only for site 1 and 2 since no allele change is observed in the contemporary population for site 3.

Additional file 2 — Supplemental tables.

Supplemental tables

**Table 7.** Elapsed time (seconds, average ± standard error (SE)) for TBLUP and SNP-BLUP approaches across scenarios. Scenarios are: all mutations having an effect (ALL), few mutations having an effect (FEW), all mutations having different effects (DIFF), the same type of mutations have the same effect (SAME), and added symmetry to SAME for reverse mutations (SYM).

| Mutation scenario | Phenotyping scenario | Approach | Average | SE |
| --- | --- | --- | --- | --- |
| ALL-DIFF | 70 | TBLUP | 2.7 | 0.1 |
|  |  | SNP-BLUP | 9.3 | 0.4 |
|  | 417 | TBLUP | 2.7 | 0.1 |
|  |  | SNP-BLUP | 27.0 | 1.0 |
|  | 1669 | TBLUP | 2.8 | 0.1 |
|  |  | SNP-BLUP | 423.0 | 159.0 |
| ALL-SAME | 70 | TBLUP | 2.6 | 0.1 |
|  |  | SNP-BLUP | 7.5 | 0.3 |
|  | 417 | TBLUP | 2.7 | 0.2 |
|  |  | SNP-BLUP | 24.8 | 1.2 |
|  | 1669 | TBLUP | 2.8 | 0.1 |
|  |  | SNP-BLUP | 245.0 | 13.8 |
| ALL-SYM | 70 | TBLUP | 2.7 | 0.1 |
|  |  | SNP-BLUP | 9.2 | 0.5 |
|  | 417 | TBLUP | 2.8 | 0.2 |
|  |  | SNP-BLUP | 27.4 | 1.1 |
|  | 1669 | TBLUP | 2.8 | 0.1 |
|  |  | SNP-BLUP | 325.0 | 16.4 |
| FEW-DIFF | 70 | TBLUP | 2.7 | 0.1 |
|  |  | SNP-BLUP | 9.5 | 0.4 |
|  | 417 | TBLUP | 2.9 | 0.2 |
|  |  | SNP-BLUP | 29.9 | 1.2 |
|  | 1669 | TBLUP | 2.9 | 0.1 |
|  |  | SNP-BLUP | 359.0 | 14.3 |
| FEW-SAME | 70 | TBLUP | 2.7 | 0.1 |
|  |  | SNP-BLUP | 8.9 | 0.5 |
|  | 417 | TBLUP | 2.8 | 0.2 |
|  |  | SNP-BLUP | 27.7 | 1.3 |
|  | 1669 | TBLUP | 2.8 | 0.1 |
|  |  | SNP-BLUP | 328.0 | 17.3 |
| FEW-SYM | 70 | TBLUP | 2.7 | 0.1 |
|  |  | SNP-BLUP | 9.7 | 0.4 |
|  | 417 | TBLUP | 2.8 | 0.2 |
|  |  | SNP-BLUP | 31.2 | 1.6 |
|  | 1669 | TBLUP | 2.9 | 0.1 |
|  |  | SNP-BLUP | 363.0 | 16.2 |

